# Phosphorylation of multiple proteins involved in ciliogenesis by Tau Tubulin kinase 2

**DOI:** 10.1101/676338

**Authors:** Ondrej Bernatik, Petra Pejskova, David Vyslouzil, Katerina Hanakova, Zbynek Zdrahal, Lukas Cajanek

## Abstract

Primary cilia (PC) are organelles necessary for proper implementation of developmental and homeostasis processes. To initiate their assembly, coordinated actions of multiple proteins are needed. Tau tubulin kinase 2 (TTBK2) is a key player in the cilium assembly pathway, controlling final step of cilia initiation. The function of TTBK2 in ciliogenesisis is critically dependent on its kinase activity, however, precise mechanism of TTBK2 action is so far incompletely understood, due to very limited information about its relevant substrates. In this study we identify CEP83, CEP89, CCDC92, Rabin8 and DVL3 as substrates of TTBK2 kinase activity. Further, we characterise a set of phosphosites of the newly identified substrates and CEP164, induced by TTBK2 *in vitro* and *in vivo*. Intriguingly, we further show that identified TTBK2 phosphosites and consensus sequence delineated from those are distinct from motifs previously assigned to TTBK2. Finally, we address functional relevance of selected phosphorylations of CEP164 and provide evidence that the examined TTBK2-induced phosphorylations of CEP164 are relevant for the process of cilia formation. In summary, our work provides important insight into substrates-TTBK2 kinase relationship and suggests that phosphorylation of substrates on multiple sites by TTBK2 is probably involved in the control of ciliogenesis in human cells.

## Introduction

Primary cilium (PC) is an organelle fundamental for proper development and homeostasis. Malfunctioning of PC leads to ciliopathies, diseases whose phenotypic manifestations are as varied as obesity, polycystic kidney disease or polydactyly (1, 2). In addition, cilia abnormalities have been recently related to cancer (3–5), further affirming the importance of correctly functioning PC for tissue homeostasis.

PC is usually formed in G1 phase of cell cycle at distal end of mother centriole, the older of the pair of centrioles which cell has at that stage. Mother centriole is distinguished from daughter centriole by two sets of proteinaceous appendages named according to their position distal (DA) and sub-distal (SDA)(6). The earliest sign of ciliogenesis seems to be accumulation of small Golgi-derived vesicles in the vicinity of DA (7–9), dependent on activity of Rab11-Rab8a-Rabin8 cascade(10–13). The vesicle docking strictly relies on intact DA, their disruption prevents the docking and in turn leads to a failure of cilium assembly (10, 14–16). Initially, five DA proteins, CEP164(17), CEP83(18), CEP89(19), FBF1(15, 20) and SCLT1(15) have been identified. Recent identification of new DA components suggests the list might not yet be complete(21, 22).

Cilia initiation turns to cilia extension process when centriolar distal end capping proteins CP110, CEP97 and MPP9 are removed and ciliary axoneme begins to elongate(23, 24). This key step in the cilium assembly pathway is controlled by Tau Tubulin kinase 2 (TTBK2)(25, 26). TTBK2 is recruited to DA by CEP164(26, 27), their mutual interaction is finetuned by INNPP5E and PI3Kγ(28). In addition, TTBK2 seems uniquely positioned among the DA-associated proteins - its depletion still allows cilia initiation to proceed to its very final stages unperturbed (vesicles dock to DA) but ciliogenesis fails at the stage when axoneme should start to extend, implying that TTBK2 may function as a switch in cilia assembly pathway, turning cilia initiation programme to cilia elongation and maintenance (25–27, 29). From evolutionary perspective, TTBK2 is a member of casein kinase 1 (CK1) superfamily, its kinase domain shows 38% identity with CK1δ(30, 31). Apart from TTBK2, additional kinase MARK4 has been implicated in cilium initiation and assembly, although it seems to control the processes via SDA rather than DA of mother centriole (32).

TTBK2 mutations leading to truncated proteins are causative for development of neurodegenerative disorder spinocerebellar ataxia type 11 (SCA11)(33–35). Interestingly, these mutant variants of TTBK2, typically truncated around 450 AA, are unable to interact with CEP164 and fail to localize to DA (25, 26), instead seem to act in dominant negative manner to disturb PC formation (25, 29). As no other kinase can compensate for defects caused by TTBK2 mutations in SCA11, it again argues that TTBK2 has non-redundant functions. Previous work established that action of TTBK2 on ciliogenesis critically depends on its kinase activity(25). However, information about *bona fide* substrates and/or S/T sites that are subjected to TTBK2 phosphorylation is very limited. So far, TTBK2 has been demonstrated to phosphorylate five substrates-Tau(36, 37), KIF2A(38), CEP164(26), CEP97(27) and MPP9(39) and only eleven S/T sites - S208 and S210 of Tau, S135 of KIF2A, S629, S636 of MPP9 and T1309, S1317, S1346, S1347, S1434 and S1443 of CEP164 (27, 38–40).

Here, we report five novel TTBK2 substrates and extensive set of TTBK2-induced phosphorylation sites determined and verified using a combination of *in vitro, in vivo* and *in silico* approaches. Using our datasets of phosphorylated peptide sequences, we delineate a kinase motif for TTBK2. Finally, we provide evidence that phosphorylation by TTBK2 modulates function of one of its targets, CEP164.

## Material and methods

### Cell culture and transfection

RPE-1 were grown in DMEM F12 (ThermoFisher Scientific, 11320033) supplemented by 10 % FBS, 1% Penicillin/Streptomycin and 1% L-glutamine, HEK293T cells were grown in DMEM Glutamax^®^ (ThermoFisher Scientific, 10569069) supplemented by 10 % FBS and 1% Penicillin/Streptomycin. Transfection of RPE1 was carried out by Lipofectamine 3000 (Invitrogen) according to manufacturer’s instructions (up to 0.5 μg DNA per condition/well, 24-well plate format). HEK293T were transfected by Polyethyleneimine (PEI) as follows: PEI was incubated in DMEM for 10min, and then mixed with plasmid equilibrated in DMEM in ratio 3 μl PEI/1μg plasmid, the mixture was incubated for 15min and then added to cells. Growth medium was changed 4h after transfection. To induce formation of PC, 24h following the transfection, RPE-1 cells were starved by serum-free medium for 24h to assess ciliogenesis or ON for TTBK2 removal measurement.

### Western blot

To analyse electrophoretic migration of proteins after TTBK2 co-expression, HEK293T cells were transfected by indicated plasmids, 24h post transfection directly lysed in 1x sample buffer (2% SDS, 62.5mM Tris pH 6.8, 10% glycerol, 2% 2-Mercaptoethanol, 0.01 % bromophenol blue). Western blot was carried out by using Bio-Rad apparatus (Mini-PROTEAN^®^ Tetra Vertical Electrophoresis Cell, Mini Trans-Blot^®^ Module) for SDS-PAGE run and transfer. Samples were loaded to discontinuous SDS-PAGE (5% stacking, 8 or 10% running gel, percentage used based on analysed protein MW), run at 150 V. Proteins were transferred to immobilon PVDF (Merck, IPVH20200) membranes at 100 V. ELFO Running buffer (0.025 M Tris, 0.192 M glycine, 0.1% SDS), WB Transfer buffer (0.025 M Tris, 0.192 M glycine, 20% Methanol). Membranes were blocked in 5% skimmed milk in TBS-Tween (20 mM Tris, 150 mM NaCl, 0.1% Tween 20, pH 7.6) and incubated with primary antibodies at 4°C ON, washed in TBS-Tween twice, incubated with secondary antibodies for 1h at RT, washed by TBS-Tween twice and developed using ECL prime (Merck, GERPN2232). For concentration of antibodies used please refer to suppl. table 3.

### Immunoprecipitation

To analyse binding of TTBK2 to CEP164 N term or CEP164 N-A term, 10 cm plates of HEK293T cells were transfected (in total 6μg of DNA mixed with 18 μl of PEI in plain DMEM). Cells were PBS washed 24h post transfection, scraped to Lysis buffer (20 mM Tris·Cl, pH 7.4, 150 mM NaCl, 0.5% NP40, 0.5% Triton-X-100, (Merck, X100), and 1× Complete proteasome inhibitors (Roche, 4693132001)) and lysed (15min, on ice). Following centrifugation (15,000 × g for 10min at +4 °C), cleared extracts were incubated ON at +4 °C in an orbital shaker with anti-FLAG sepharose beads (M2 clone, Merck, A2220). Bound complexes were pelleted, washed, and analysed by SDS-PAGE and Western blotting. Where appropriate, contrast and/or brightness of images were adjusted by using Photoshop CS5 (Adobe).

### Purification of proteins from HEK293T cells

Cep164 N-, M- and C-terminal fragment, CEP83, CEP89, CCDC92, Rabin8, DVL3 coding plasmids fused to FLAG tag and FLAG-TTBK2 WT and kinase dead variant (KD) were used for production of proteins for *in vitro* or TTBK2 autophosphorylation experiments, GFP-TTBK2 WT was coexpressed for *in vivo* experiments. To purify protein of interest, each expression plasmid (please see detailed list of used plasmids in suppl. table 3) was transfected to HEK293T cells (2× 15 cm plate per condition, 20 μg DNA), either individually (*in vitro* kinase assay) or in combination with TTBK2 expression vector (*in vivo*). 36h post transfection the proteins of interest were immunoprecipitated as follows: cells were washed in PBS, scraped, pelleted by centrifugation (400g/ 5min/ +4°C), resuspended in Lysis buffer (20 mM Tris·Cl, pH 7.4, 150 mM NaCl, 0.5% NP40, 0.5% Triton-X-100, 0.1 mM DTT (Merck, 43815), 1× Complete proteasome inhibitors (Merck, 4693132001), 1x phosphoSTOP (Merck, PHOSS-RO), sonicated (2×20sec, on ice), and lysed (10min, on ice). Following centrifugation (15,000 × g for 10min at +4°C), cleared extracts were incubated 2h-6h at +4°C in an orbital shaker with anti-FLAG sepharose beads (M2 clone, Merck, A2220). Bound complexes were pelleted and 5x washed (Lysis buffer with 1M NaCl). In case of samples for *in vitro* kinase assay, those were in addition 2x washed with kinase buffer (50 mM Tris-Cl ph=7.4, 10mM MgCl2, 0.1 M EGTA) +0.1 mM DTT, resuspended in kinase buffer + 0.1% DTT + 10% glycerol and stored as 25% slurry at −80 °C until used. Where indicated, samples were treated with λ-phosphatase (New England BioLabs, P0753S) (30 °C for 20min) according to manufacturer’s instructions, λ-phosphatase was then removed by 2x wash with kinase buffer + phosphoSTOP (Merck, PHOSS-RO), 50mM NaF (NEB, P0759S) and 10mM sodium orthovanadate (NEB, P0758S). Samples were subsequently subjected to *in vitro* kinase assay (*in vitro* samples) or directly analysed by MS/MS (*in vivo* samples).

### Immunocytofluorescence microscopy

Cells seeded on glass coverslips were fixed for 10min in −20°C Methanol, washed 3x with PBS, blocked (2% BSA in PBS with 0.01% NaN3), 3x PBS washed, incubated with primary antibodies for 1h, 3x washed by PBS, incubated with secondary antibodies for 1h in dark, washed 3x with PBS, incubated with DAPI, washed by PBS 2x and mounted by glycergel (DAKO #C0563). Microscopy analyses were done using either DeltaVision Elite (GE Healthcare) with 100x/Zeiss Plan-ApoChromat 1.4 objective and DeltaVision softWoRx acquisition SW (here the image stacks were taken with a Z distance of 0.2 μm, deconvolved with one cycle of conservative ratio deconvolution algorithm, and directly projected as maximal intensity images) or Zeiss AxioImager.Z2 with Hamamatsu ORCA Flash 4.0 camera, 100x Apo oil immersion objective, and ZEN Blue 2.6 acquisition SW (Zeiss). Image stacks acquired using Zeiss AxioImager.Z2 were projected as maximal intensity images by using ImageJ distribution FIJI(41). Where appropriate, contrast and/or brightness of images were adjusted by using Photoshop CS5 (Adobe) or FIJI. For ciliogenesis experiment (Fig. 4C), at least 50 cells were analysed per each condition. For experiment in Fig. 4D, a densitometry analysis within selected regions of interest was performed in 16-bit TIFF images by using FIJI, analysing 30-40 cells per condition. Data are presented as relative staining intensity (staining intensity of a protein of interest normalized to the staining intensity of the centriolar marker CAP350). Statistical analysis by unpaired Student t-test were performed using Graphpad Prism, P<0.05 (*), P<0.01 (**), P<0.001 (***) and P<0.0001 (****) were considered significant. Results are presented as mean plus SEM.

**Figure 1:**
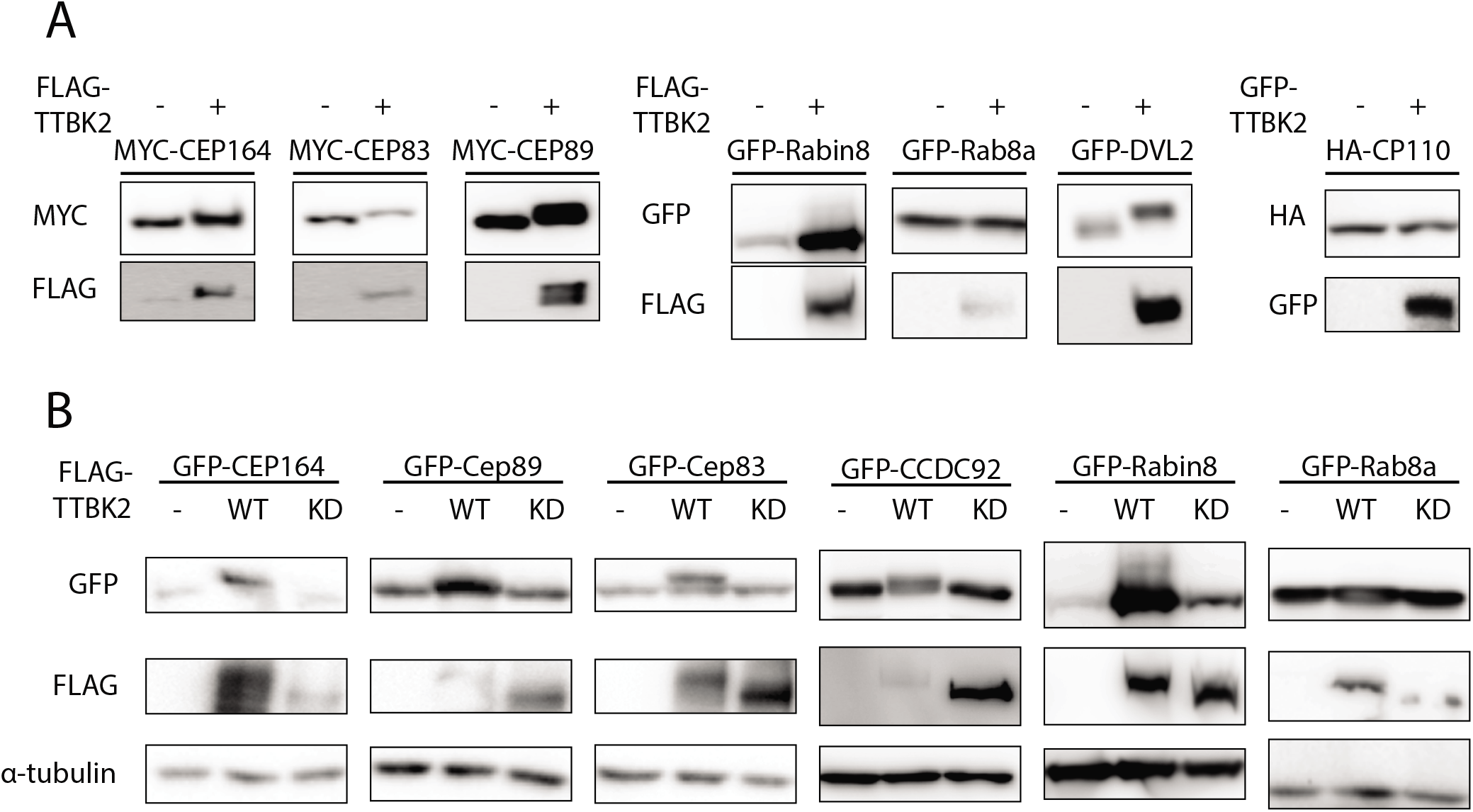
TTBK2 induces mobility shift of of CEP164, CEP89, CEP83, CCDC92, Rabin8 and DVL3. HEK293T cells were transfected by indicated plasmids, lysed 24h post-transfection and analysed by WB using indicated antibodies. **A:** Electrophoretic mobility of MYC-CEP164, MYC-CEP83, MYC-CEP89, GFP-Rabin8 and GFP-DVL2 is decelerated by FLAG-TTBK2 expression. CP110 and Rab8a do not change their mobility upon FLAG-TTBK2 or GFP-TTBK2 expression. **B:** Detection of mobility shift of GFP-CEP164, GFP-CEP89, GFP-CEP83, GFP-CCDC92, and GFP-Rabin8, induced by FLAG-TTBK2 but not FLAG-TTBK2 kinase dead (KD). Note that TTBK2 WT undergoes profound mobility shift compared to TTBK2 KD, indicating autophoshorylation.

**Figure 2:**
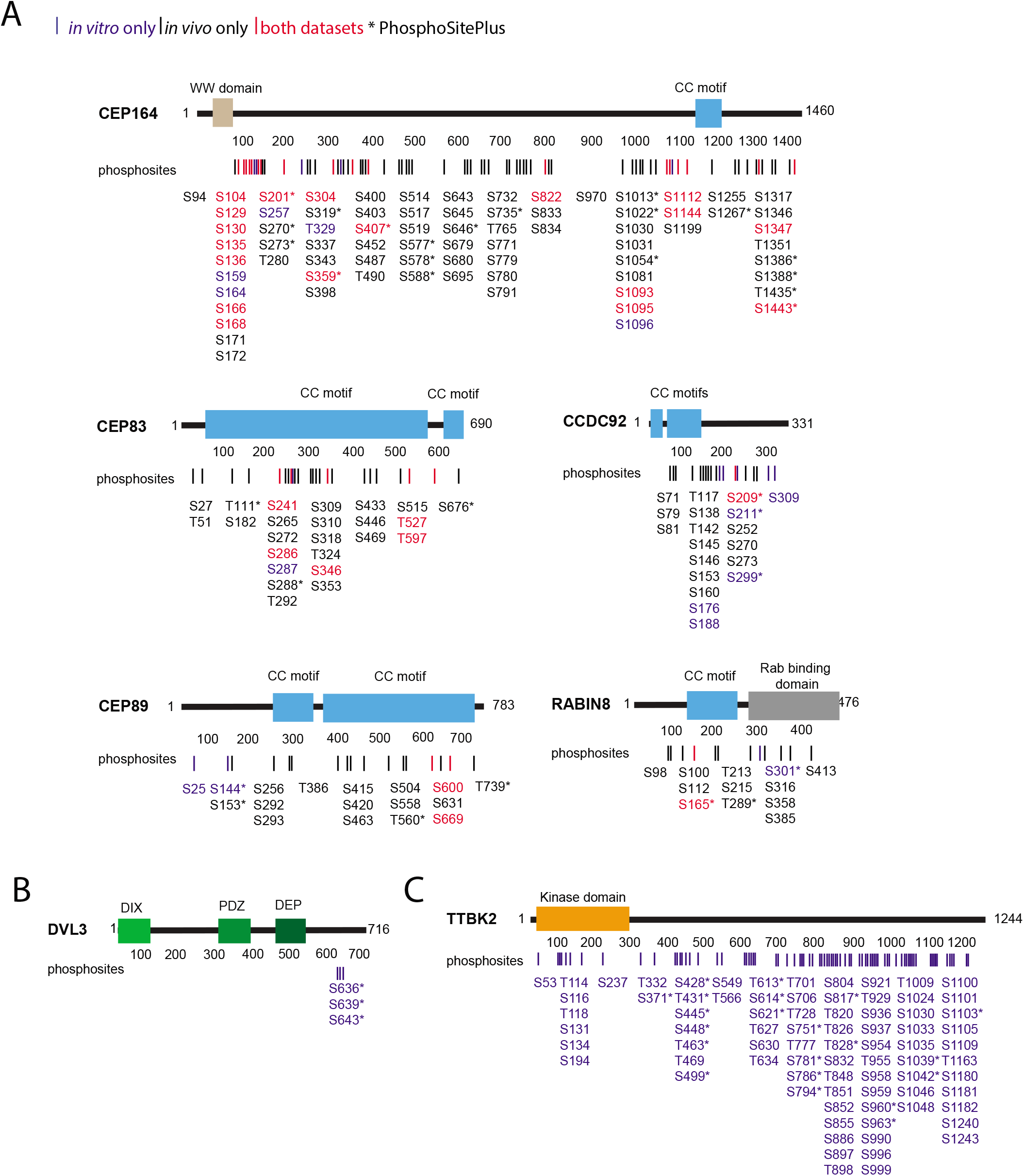
TTBK2 phosphorylates its substrates on multiple serines and threonines. Map of identified phosphorylations induced by TTBK2 on tested substrates. **A, B:** Structure of each TTBK2 substrate is schematized, rectangles indicate presence of a domain or motif, and numbers indicate length of given protein in amino acids. Lines and numbers, respectively, below the schematic protein structure indicate positions of each phosphorylation induced by TTBK2. Sites shown in red were detected both in vitro and in vivo, sites in blue were detected only in vitro and sites in black only in vivo. Asterisk indicates that given phosphorylation is covered in PhosphoSitePlus. **A:** TTBK2 induced phosphorylation of CEP164, CEP89, CEP83, CCDC92 or Rabin8 identified in vitro and in vivo **B:** TTBK2 phosphorylation of DVL3 identified in vitro **C:** TTBK2 autophosphorylation identified in vitro.

**Figure 3:**
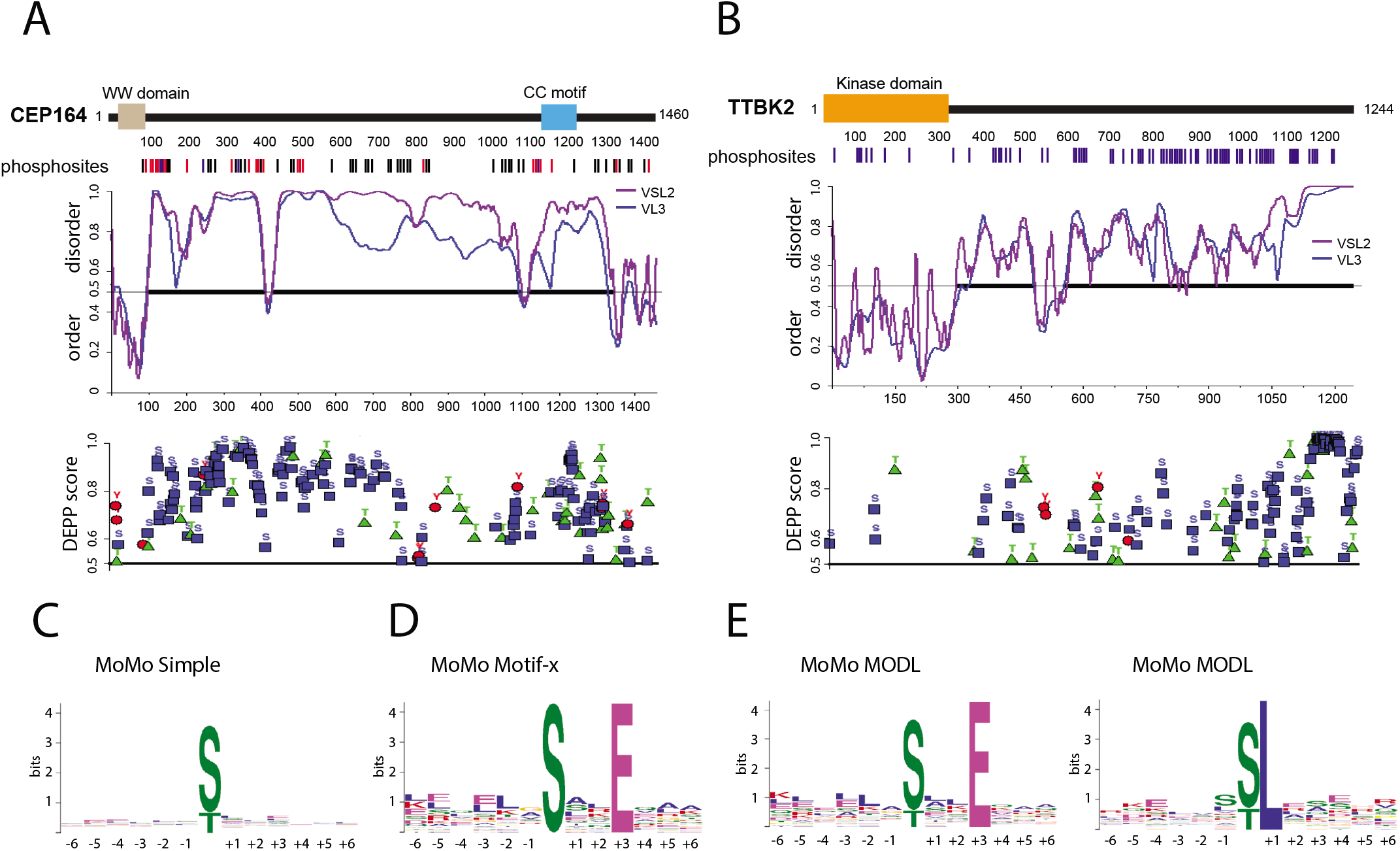
TTBK2 shows motif similarities to CK1. **A, B:** Protein disorder and phosphorylation prediction for CEP164 and TTBK2. Structure of CEP164 and TTBK2 is schematized as described in Fig. 2. The first graph below schematized structure of CEP164 and TTBK2, respectively, depicts prediction of disorder/order (with cut-off 0.5) determined by PONDR using VSL2 (Violet line) and VL3-BA (Blue line) algorithms. The second graph plots Disorder Enhanced Phosphorylation Predictor (DEPP) score (with cut-off score 0.5) and individual phosphosites thereby predicted (pS as blue rectangle, pT as green triangle, and phosphoY as red circle). **C, D, E:** Determination of TTBK2 kinase motifs using indicated algorithms. Size of depicted symbols indicates relative abundance of each aminoacid at given position within a set of phosphorylated peptides. Kinase motif search was performed by **C:** MoMo simple, **D:** Motif-x, and **E:** MODL alogrithms.

**Figure 4:**
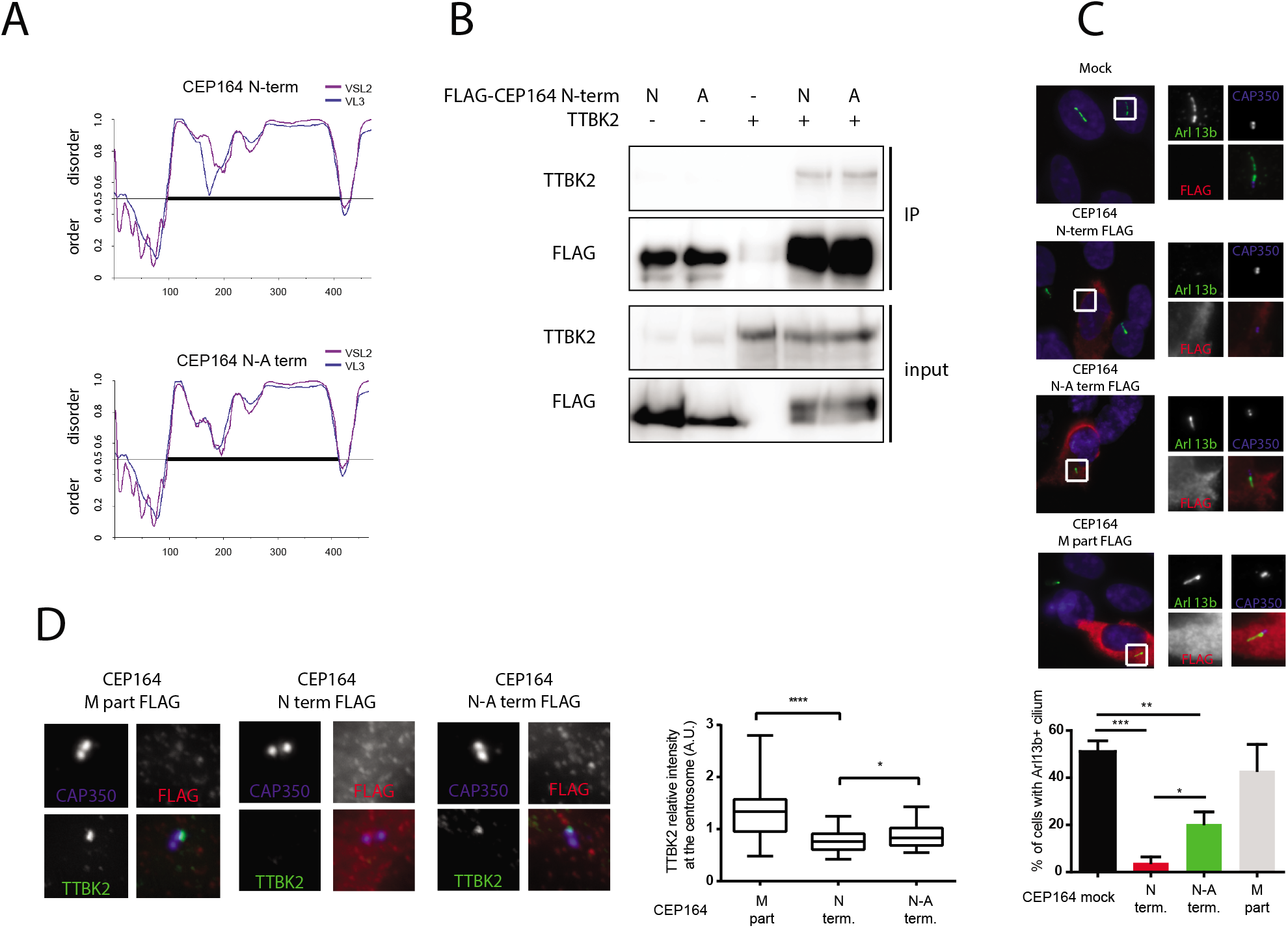
Phosphorylation of CEP164 N-terminus by TTBK2 affects its function and ciliogenesis. **A:** In silico analysis of protein folding by PONDR VSL2 (Violet line) and VL3-BA (Blue line) algorithms, the numbers indicate length of given protein in amino acids. Note that mutation of seven TTBK2 phosphorylated sites (S129, S130, S135, S136, S166, S168, S201) to Alanines (CEP164 N-A term) is not expected to change protein folding over control (CEP164 N-term). **B:** HEK293T cells were transfected by indicated plasmids and 24h post-transfection were subjected to immunoprecipitation by anti-FLAG antibody. No difference was observed between FLAG-CEP164 N term and FLAG-CEP164 N-A term in their ability of to co-immunoprecipitate MYC-TTBK2. **C-D:** RPE1 cells were transfected by indicated FLAG-tagged CEP164 truncation constructs and, following 24h, starved for additional 24h to induce ciliogenesis or ON to evaluate TTBK2 sequestration. **C:** ARL13b (green) and CAP350 (blue, inset) staining was used to detect PCs and centrioles, respectively, expression of indicated CEP164 variants was detected by FLAG antibody (red). The graph summarizes effects on PC formation from 4 independent experiments, RPE-1 cells transfected with empty vector (mock) or CEP164 M part were used as controls. Analysed by student’s t-test *** p<0.001, ** p<0.01, * p<0.05, SEM. **D:** Following fixation, RPE-1 cells were stained for TTBK2 (green), FLAG (red), and CAP350 (blue) to examine differences in localization of endogenous TTBK2 to mother centriole. Graph summarizes the effects on TTBK2 sequestration (n=5, 30-40 cells analysed per condition). Analysed by student’s t-test, **** p<0.0001, * p<0.05, SEM.

### In vitro kinase assay

Beads-bound proteins were washed with kinase buffer (50 mM Tris, pH 7.4, 10 mM MgCl2, and 0.1 mM EGTA), mixed according to experimental scheme (Supp. Fig. 1) and incubated in the kinase buffer supplemented with 1 mM ATP, 5x Complete proteasome inhibitors (Merck, 4693132001), 1x phosphoSTOP (Merck, PHOSS-RO), 50mM NaF (NEB, P0759S) and 10mM sodium orthovanadate (NEB, P0758S), and incubated for 30 minutes/30°C. The kinase reaction was terminated by addition of 4x sample buffer, and the resulting samples were analysed by MS/MS.

### MS/MS analysis

#### In gel digestion

Immunoprecipitates were separated on SDS-PAGE gel electrophoresis, fixed with acetic acid in methanol and stained with Coomassie brilliant blue G250 (EZBlue, G1041, Merck) for 1 hour. Corresponding 1D bands were excised. After destaining, the proteins in gel pieces were incubated with 10 mM DTT at 56 °C for 45 min. After removal of DTT excess, samples were incubated with 55 mM IAA at room temperature in darkness for 30 min, then alkylation solution was removed and gel pieces were hydrated for 45 min at 4°C in digestion solution (5 ng/μl trypsin, sequencing grade, Promega, in 25 mM AB). The trypsin digestion proceeded for 2 hours at 37°C on Thermomixer (750 rpm; Eppendorf). Digested peptides were extracted from gels using 50% ACN solution with 2.5% formic acid (FA) and concentrated in speedVac concentrator (Eppendorf). The aliquot (1/10) of concentrated sample was transferred to LC-MS vial with already added polyethylene glycol (PEG; final concentration 0.001%, (49)) and directly analyzed by LC-MS/MS for protein identification.

#### Phosphopeptide enrichment

The rest of the sample (9/10) was used for phosphopeptide analysis. Sample was diluted with acidified acetonitrile solution (80% ACN, 2% FA). Phosphopeptides were enriched using Pierce Magnetic Titanium Dioxide Phosphopeptide Enrichment Kit (Thermo Scientific, Waltham, Massachusetts, USA) according to manufacturer protocol and eluted into LC-MS vial with already added PEG (final concentration 0.001%). Eluates were concentrated under vacuum and then dissolved in water and 0.6 μl of 5% FA to get 12 μl of peptide solution before LC-MS/MS analysis.

#### LC-MS/MS analysis

LC-MS/MS analyses of peptide mixture were done using RSLCnano system connected to Orbitrap Elite hybrid spectrometer (Thermo Fisher Scientific) with ABIRD (Active Background Ion Reduction Device; ESI Source Solutions) and Digital PicoView 550 (New Objective) ion source (tip rinsing by 50% acetonitrile with 0.1% formic acid) installed. Prior to LC separation, peptide samples were online concentrated and desalted using trapping column (100 μm × 30 mm) filled with 3.5 μm X-Bridge BEH 130 C18 sorbent (Waters). After washing of trapping column with 0.1% FA, the peptides were eluted (flow rate 300 nl/min) from the trapping column onto Acclaim Pepmap100 C18 column (3 μm particles, 75 μm × 500 mm; Thermo Fisher Scientific) by 65 min long gradient. Mobile phase A (0.1% FA in water) and mobile phase B (0.1% FA in 80% acetonitrile) were used. The gradient elution started at 1% of mobile phase B and increased from 1% to 56% during the first 50 min (30% in the 35^th^ and 56% in 50^th^ min), then increased linearly to 80% of mobile phase B in the next 5 min and remained at this state for the next 10 min. Equilibration of the trapping column and the column was done prior to sample injection to sample loop. The analytical column outlet was directly connected to the Digital PicoView 550 ion source. MS data were acquired in a data-dependent strategy selecting up to top 10 precursors based on precursor abundance in the survey scan (350–2000 m/z). The resolution of the survey scan was 60 000 (400 m/z) with a target value of 1 × 10^6^ ions, one microscan and maximum injection time of 1000 ms. High resolution (15 000 at 400 m/z) HCD MS/MS spectra were acquired with a target value of 50 000. Normalized collision energy was 32% for HCD spectra. The maximum injection time for MS/MS was 500 ms. Dynamic exclusion was enabled for 45 s after one MS/MS spectra acquisition and early expiration was disabled. The isolation window for MS/MS fragmentation was set to 2 m/z.

#### Data analysis

The analysis of the mass spectrometric RAW data was carried out using the Proteome Discoverer software (Thermo Fisher Scientific; version 1.4) with in-house Mascot (Matrixscience; version 2.4.1 or higher) search engine utilization. MS/MS ion searches were done against in-house database containing expected protein of interest with additional sequences from cRAP database (downloaded from http://www.thegpm.org/crap/). Mass tolerance for peptides and MS/MS fragments were 7 ppm and 0.03 Da, respectively. Oxidation of methionine, deamidation (N, Q) and phosphorylation (S, T, Y) as optional modification, carbamidomethylation of C as fixed modification, no enzyme specifity was selected. The phosphoRS (version 3.1) feature was used for preliminary phosphorylation localization. Final phosphosite assignment (including inspection of phosphorylations with ambiguous localization) was performed by manual evaluation of the fragmentation spectra of the individual phosphopeptides. Quantitative information was assessed and manually validated in Skyline software (Skyline daily 3.6.1.10230). Normalization of the data was performed using a set of phosphopeptide standards (added to the sample prior phosphoenrichment step; MS PhosphoMix 1, 2, 3 Light, Sigma) in combination with summed peak area of non-phosphorylated peptides identified in analyses of samples without phosphoenrichment. The mass spectrometry proteomics data have been deposited to the ProteomeXchange Consortium via the PRIDE(42) partner repository with the dataset identifier PXD014199.

### Identification of phosphorylations

Average intensity for individual phosphosites or clusters was calculated from MS/MS data as a sum of intensities of all phosphorylated peptides containing the particular pS/pT. Data are summarized in Suppl. table 1 and graphically represented in Suppl. Fig. 2A, Suppl. Fig. 2B, Suppl. Fig. 3. Induction of phosphosites by TTBK2, summarized for individual substrates in Fig.2, was evaluated using individual phosphorylated peptides intensities to allow careful curation of the data. We used following criteria to assess induction of phosphosite by TTBK2:

*in vitro* dataset. 1: intensity of at least one phosphorylated peptide of given phosphosite must have been equal to or higher than 2×10^6^ threshold. 2: sum of intensities of all phosphopeptides detected for given phosphosite in individual experiment must have been at least 2-fold of total intensity of the corresponding phosphosite in control sample (substrate incubated with TTBK2 kinase dead (KD)). If exact position of phosphorylated residue could not be resolved, each of S/T residues of given cluster was considered as phosphorylated. If phosphopeptide containing such cluster met criterium 1 and 2, the cluster was considered induced (in total 6 peptides with 15 individual phosphorylations).

In case of TTBK2 autophosphorylation analysis, two reference samples (controls) were used (②TTBK2 WT + λ-phosphatase; ④TTBK2 KD + λ-phosphatase + kinase assay), as shown in Suppl. Fig. 1B. Thus, the individual phosphorylation must have fulfilled the above described criteria for both controls to be considered TTBK2 induced.

*in vivo* dataset. 1: intensity of at least one phosphorylated peptide must have been equal to or higher than 2×10^6^ threshold if identified repeatedly, or equal to or higher than 2×10^7^ threshold if detected in a single experiment. 2: sum of intensities of all phosphopeptides detected for given phosphosite in individual experiment must have been at least 2-fold of intensity of the corresponding phosphosite in control sample (Mock). If exact position of phosphorylated residue could not be resolved, each of S/T residues of given cluster was considered as phosphorylated. If phosphopeptide containing such cluster met criterium 1 and 2, cluster was considered induced (in total 2 peptides with 4 individual phosphorylations).

Graphs in Suppl. Fig. 2A, Suppl. Fig. 2B, Suppl. Fig. 3 show comparison of average intensity for individual S/T phosphosites. Cases where the exact position could not be assigned are indicated as (/) for two possible positions or (-) for clusters. **Supplementary Fig. 2A:** For every S/T phosphosite average intensity detected in samples with Mock (Black bar), TTBK2 WT induced (Red bar), Mock + λ phosphatase (Grey bar) and TTBK2 WT + λ phosphatase (Blue bar) is indicated on y axis. To allow better visualisation of intensity of individual phosphosites in one graph, note that the scale of y axis is in log10. **Supplementary Fig. 2B:** For every S/T phosphosite average intensity detected in samples with ①TTBK2 WT (Black bar), ② CTRL#1: TTBK2 WT + λ phosphatase (Grey bar), ③TTBK2 WT + λ phosphatase + kinase assay (Red bar), and ④ CTRL#2: TTBK2 KD + λ phosphatase + kinase assay (Brown bar) is indicated on y axis, log10 scale. **Supplementary Fig. 3:** For every S/T phosphosite average intensity detected in samples with Mock (Black bar), TTBK2 WT induced (Red bar), is indicated on y axis, log10 scale.

### Kinase motif determination

Kinase motifs were delineated using dataset of 13 AA long peptides with central S/T, which we identified as TTBK2-induced. For the purpose of this analysis, the *in vitro* and *in vivo* datasets were cross-correlated and phosphorylations identified as TTBK2 induced in one dataset and not induced in the other were excluded from kinase motif determination peptide dataset. Peptide sequences was loaded to MeMe suite phosphorylation motif finder MoMo, and analysed using simple, Motif-x and MODL algorithms(43, 44). Minimal number of occurrences for kinase motif to be considered was set to 10 for all algorithms. Specific setting for Motif-x algorithms was p value <0.001, motifs with central S/T were considered separately. Default MODL settings were used (Maximum number of iterations = 50, Maximum number of iterations with no decrease in MDL=10, with central S/T combined onto one motif). Peptides used and those identified by individual algorithms are listed in Suppl. Table 2.

### In silico analyses

Depiction of domains or motifs in Fig. 2A, Fig. 2B, Fig. 2C, Fig. 3A, 3B, Suppl. Fig. 4, Suppl. Fig. 5 was based on Uniprot.org. Canonical proteins sequences from uniprot.org were used for disorder and phosphorylation prediction (Fig. 3A, Fig. 3B, Suppl. Fig. 4, Suppl. Fig. 5), protein identifiers used are as follows: CEP164 (<u>Q9UPV0</u>). CEP83 (<u>Q9Y592</u>), CEP89 (<u>Q96ST8</u>). CCDC92 (<u>Q53HC0</u>), Rabin8 (<u>Q96QF0</u>), DVL3 (<u>Q92997</u>), TTBK2 (<u>Q6IQ55</u>), MPP9 (<u>Q99550</u>), KIF2a (<u>O00139</u>), Tau (<u>P10636</u>), CEP97 (<u>Q8IW35</u>). Prediction of protein disorder was performed by Predictor of Natural Disordered Regions (PONDR) (http://www.pondr.com/) algorithms VSL-2(45) and VL3-BA(46). Score above 0.5 for disorder prediction suggests unstable or none secondary structure, score below 0.5 suggest this protein part has defined secondary structure. Prediction of protein phosphorylation was performed by the Disorder Enhanced Phosphorylation Predictor (DEPP)(47) (http://www.pondr.com/cgi-bin/depp.cgi). Only phosphorylations with DEPP score above 0.5 were considered as possibly phosphorylated, with the highest scoring phosphorylations being the most probable.

## Results

### TTBK2 acts on multiple candidate substrates

To identify novel TTBK2 substrates, we focused our effort on a set of candidate proteins that localize to DA or interact with CEP164 and have been implicated in cilia initiation. These include DA components CEP83 and CEP89, Rab8a, Rabin8, DVL, CCDC92 and CP110 (10, 15, 24, 48, 49). First, given that reported TTBK2 substrates CEP164 or MPP9 respond to elevated TTBK2 levels by profound mobility shift (26, 39) we examined behaviour of individual candidates following their co-expression with TTBK2 in HEK293T cells. Interestingly, while we found no response to FLAG-TTBK2 co-expression in case of HA-CP110 and GFP-Rab8a, we observed a mobility shift of MYC-CEP83, MYC-CEP89, GFP-Rabin8, and GFP-DVL2 similar to that of MYC-CEP164 (Fig. 1A), indicating that these proteins are modified in TTBK2-dependent manner. Next, to corroborate our observations, we included TTBK2 D163A mutant with disabled kinase domain activity (KD)(50) in our experiments. Importantly, as depicted in Fig. 1B, GFP-CEP164, GFP-CEP89, GFP-CEP83, GFP-CCDC92, and GFP-Rabin8, but not GFP-Rab8a, specifically responded to TTBK2, but not TTBK2 KD, suggesting the observed shifts in protein mobility reflect modification of given protein mediated by kinase activity of TTBK2.

### TTBK2 directly phosphorylates CEP83, CEP89, Rabin8, CCDC92 and DVL3

Our initial results implied that TTBK2 might phosphorylate several proteins necessary for the process of cilia initiation. To confirm and extend this hypothesis, we performed *in vitro* kinase assays of individual candidate substrates in the presence of TTBK2, followed by MS/MS identification and quantification of detected phosphorylations. To ensure proper protein folding and functionality, individual candidate substrates and TTBK2, respectively, were purified by immunoprecipitation from HEK293T cells. TTBK2 KD was used as a control condition to account for actions of endogenous kinases possibly co-purifying with immunoprecipitated TTBK2. In addition, in the second repetition we included λ phosphatase treatment into our workflow to reduce impact of any baseline phosphorylation already present in the purified candidate substrates. Outline of the experimental workflow is summarized in Suppl. Fig. 1A. In short, the criteria to assign any identified phosphorylation to TTBK2 were following (please see M&M for additional details): a) intensity of the phosphopeptide was higher than 2×10^6^ threshold b) intensity for analysed phosphosite was increased to at least two folds of the corresponding intensity in control samples (TTBK2 KD). By applying these criteria, we detected 45 phosphorylated S/T sites induced by TTBK2 in the tested proteins (CEP164, CEP83, CEP89, Rabin8, CCDC92, DVL3) (Fig. 2A, Fig. 2B, blue and red colour coded). Total number of identified phosphorylations was ~120 (Suppl. Fig. 2A, Fig 2B, Suppl. Table 1).

To extend our analysis, we in similar manner examined autophosphorylation of TTBK2. To identify autophosphorylation sites, we analysed four experimental conditions: ①purified TTBK2, ②purified TTBK2 treated with λ phosphatase, ③ TTBK2 treated with λ phosphatase subsequently subjected to *in vitro* kinase assay, and ④ TTBK2 KD treated with λ phosphatase, subjected to *in vitro* kinase assay (Schematized in Suppl. Fig. 1B). We compared phosphosites identified in the condition ③ with two control conditions, the dephosphorylated TTBK2 WT without kinase assay ② and the dephosphorylated *in vitro* kinase assay of TTBK2 KD ④. Phosphorylations that were at least 2-fold induced over both controls were considered as TTBK2 induced. Condition ① was included to assess efficiency of λ phosphatase treatment. This analysis led to identification of approximately 110 phosphorylations, out of which 79 were found induced by TTBK2. The induced phosphorylations were distributed along the whole sequence of TTBK2, with significant portion of the sites residing to C-terminus (Fig. 2C, Suppl. Fig. 2B, Suppl. Table 1).

Identification of specific S/T residues phosphorylation by *in vitro* kinase assay typically suffices to assume direct kinase-substrate relationship for individual proteins tested but might not fully reflect phosphorylation of a given protein *in vivo*. To this end, we set to analyse phosphorylations of individual substrates co-expressed with TTBK2 and subsequently purified from HEK293T cells. The experiment workflow is summarized in Suppl. Fig. 1C. Criteria for TTBK2-induced phosphorylations were set more stringent than in the case of *in vitro* analysis, to account for higher complexity of the samples and in turn for potential influence of other protein kinases (please refer to M&M for details). This *in vivo* approach identified approximately 230 phosphorylations (Suppl. Table 1, Suppl. Fig. 3), out of which 133 were induced by TTBK2 (Fig. 2A, black and red colour coded). Importantly, comparison of induced phosphorylation sites identified *in vivo* and *in vitro* showed not only fairly extensive overlap between the datasets for individual proteins (phosphorylation common for both datasets are red, Fig. 2A, Suppl. Fig. 3), but also revealed that many phosphosites localized in clusters or pairs, often dispersed through the entire protein sequence (Fig. 2A).

Specifically, in case of CEP164, 18 individual S/T sites were found induced by TTBK2 by both approaches, for CEP83 it was 5 S/T residues, for CEP89 2 S/T and, and CCDC92 and Rabin both showed 1 S/T phosphorylation in common between *in vitro* and *in vivo*. Of note, analysis of DVL3 *in vivo* phosphorylations induced in context of several kinases, including TTBK2, has been recently reported (51). Together, 27 (64%) of induced phosphorylations detected by *in vitro* approach were also found induced *in vivo*, hence confirming a role of TTBK2 in phosphorylation of tested proteins (Fig. 2A). Additionally, 6 sites identified as induced *in vitro* were occasionally detected also *in vivo* but did not pass the set threshold to be considered induced (Suppl. Table 1, Suppl. Fig. 2A, Suppl. Fig. 3). Noteworthy, significant portion of sites detected by *in vivo* analyses (in total 106 phosphorylated S/T residues, Suppl. Table 1) were not detected using the *in vitro* approach.

### TTBK2 shows motif similarities to CK1

Given the number of phosphorylations we identified and the reports on preferential phosphorylation of IDRs over folded protein regions (47, 52–54) we inspected hereby analysed TTBK2 substrates by Predictor of Natural Disordered Regions (PONDR)(45, 46) and Enhanced Phosphorylation Predictor (DEPP)(47), to check the extent of IDRs and the overlap between the identified and the predicted phoshosites, respectively. The analysis carried using PONDR algorithms VSL-2 (Fig. 3A, 3B, Suppl. Fig. 4, violet line) and VL3-BA (Fig. 3A, 3B, Suppl. Fig. 4, blue line), suitable for examination of long IDRs, revealed that all proteins tested are predicted to contain long IDRs (Fig. 3A, 3B and Suppl. Fig. 4). In addition, comparing our experimental data to DEPP-predicted phosphorylations revealed many examples of extensive overlap (Fig. 3A, 3B, Suppl. Fig. 4). Specifically, experimentally identified CEP164 phosphorylations were found distributed throughout the protein sequence in a pattern resembling that of DEPP prediction (Fig. 3A). Further, DEPP predicted extensive phosphorylation of TTBK2 C-terminal part, in good agreement with our experimental data (Fig. 3A). In addition, we observed some extent of similarity between the experimentally detected and the predicted phosphorylation sites also in the case of the remaining substrates of TTBK2 we examined (Suppl. Fig. 4). Given these findings, we next examined also the previously identified substrates and phosphosites of TTBK2. As expected, all previously identified phosphosites were localized within long IDRs, which were predicted to contain many additional phosphorylation sites (Suppl. Fig. 5).

To validate our data with previously unbiasedly detected phosphorylations on analysed proteins, we cross-examined our results with PhosphoSitePlus database. For the purpose of this comparison, we did not discriminate here between induced and noninduced phosphorylations or between individual datasets of our analysis. Interestingly, we found that 99/133 (74%) phosphorylations reported for substrate proteins on PhosphoSitePlus were also detected in our experiments (Fig. 2A, 2B, 2C Suppl. Table 1, marked by asterisk). As many phosphorylations identified by us were previously detected in endogenous proteins it suggests they are not artefactual, although we cannot rule out some are a result of overexpression. Importantly however, only 53/229 (23 %) of TTBK2 induced phosphorylations were previously identified by large-scale proteomic studies. This implies that phosphorylation of some of these residues might be limited to specific cellular context.

As it was previously reported that TTBK2 preferentially phosphorylated S/T sites with pY at +2 position(50), we examined out datasets for the presence of this reported preferred kinase motif. However, we identified only 2 phosphorylations (S201 in CEP164, S316 in Rabin8) with Y at position +2, which could in principle fit into the proposed consensus sequence. Since consensus motif reported for CK1 family to which TTBK2 belongs to is pS/pT-x-x-S/T(55), we manually searched the dataset for priming sites at position −3 and found 28 phosphorylations fitting this criteria (CEP164: S171, S273, S403, S455, T490, S517, S646, S735, S1096, S1255, S1258; CCDC92: S145, S179, S273; Rabin8: S100; DVL3: S639; TTBK2: S134, T431, S448, T820, T851, S855, S958, S963, S999, S1033, S1042, S1243). Thus, majority of phosphorylation sites we found were without any kinase motif assigned. To probe for possible unrecognized TTBK2 consensus sites, we pooled data from *in vitro* and *in vivo* dataset and used them for unbiased kinase motif search by MeMe suite tool MoMo (43, 56). Initial analysis by “simple” algorithm, which plots relative abundance of individual AA residues at indicated position within the whole dataset, did not reveal any enrichment (Fig. 3C). Given that, we subsequently performed analysis by Motif-x and MODL algorithms, respectively, that can resolve consensus motif sequences present only in a subset of peptides (44, 56). Intriguingly, both algorithms detected preference for glutamic acid (E) on position +3 (MODL, 32 peptides, Motif-x, 27 peptides) (Fig. 3D, 3E). In addition, analysis using MODL detected preference for leucine (L) on position +1 (MODL, 30 peptides) (Fig. 3E). Sequences of peptides carrying found motifs are listed in the Suppl. Table.2 and their position is indicated for individual proteins analysed.

### CEP164 N-terminus phosphorylation leads to altered function

Functional relevance of a phosphorylation is usually tested by mutating detected phosphorylated sites to non-phosphorylable variants and subsequent examination of whether this alters function of tested protein (57). Our analysis of phosphorylation of several target proteins by TTBK2 revealed significant number of induced phosphorylation sites, implying possible complex regulation as multiple phosphorylations typically act in combinatorial fashion (53, 58). This presented us with a conundrum of how to test functional relevance of identified phosphorylations, firstly; as they were tens of them and secondly; as most of them resided to IDRs with currently unknown impact on biology of a given protein. Having said that, we focused our functional validation efforts on TTBK2 sites identified within N-terminal part of CEP164 (CEP164 N-term, AA 1-467), which, via its WW domain interacts with TTBK2 and hence affects ciliogenesis (10, 26, 27). We selected phosphorylated residues (S129, S130, S135, S136, S166, S168, S201) located in the proximity of WW domain and mutated these in CEP164 N-term to A (to obtain CEP164 N-A term) to examine their possible role in the WW-domain mediated interaction between CEP164 and TTBK2. Noteworthy, *in silico* modelling of introduced mutations using PONDR did not reveal any major impact on protein folding (Fig. 4A). To resolve if the introduced mutations in any way changed CEP164-TTBK2 relationship, we first tested whether CEP164 N-A term and TTBK2 are still able to co-immunoprecipitate and did not detect any notable difference between FLAG-CEP164 N-term and FLAG-CEP164 N-A term binding to MYC-TTBK2 when over-expressed in HEK293T cells (Fig. 4B). We reasoned that our failure to see any difference meant the changes were too subtle to be revealed by the used approach and decided to test this possibility by other means. Interestingly, we observed that while ciliogenesis was efficiently induced in mock (51%) or FLAG-CEP164 M-part (468-1135 AA) (43%) transfected RPE-1 cells, FLAG-CEP164 N-term expression almost completely abrogated formation of PC (below 5% of ciliated cells in agreement with previous reports (10, 26, 27). In contrast, expression of FLAG-CEP164 N-A term still allowed formation of PC in about 20% of transfected RPE-1 cells (Fig. 4C). Based on this observation we hypothesized that ability of CEP164 N-term to sequester endogenous TTBK2(26) might be affected in CEP164 N-A term. Intriguingly, we found that FLAG-CEP164 N-A term was able to sequester TTBK2 from mother centriole, but to a lesser extent that FLAG-CEP164 N-term (Fig. 4D). These results suggest that phosphorylation of given residues (S129, S130, S135, S136, S166, S168, S201) is functionally important for TTBK2 recruitment to mother centriole and hence ciliogenesis regulation.

## Discussion

TTBK2 seems to have unique position among other regulators of ciliogenesis, as its action marks “end of beginning” by pushing cilia initiation towards cilia extension. However, understanding how TTBK2 regulatory function is implemented is still only partial. To gain insight into the process, we set to identify its substrates. In this study we demonstrated that several proteins that either directly bind to CEP164, or localize to its proximity, namely CEP83, CEP89, CCDC92, Rabin8 and DVL3, and CEP164 itself, are subjected to TTBK2 phosphorylation, primarily in their IDRs. We further revealed that TTBK2 shows previously undetected consensus motif similarity to that of CK1, at least for a subset of phosphorylated sites.

Our observation that TTBK2 is able to phosphorylate components of DA and proteins associated with centriole distal end does not come as a complete surprise, given its proposed role in the assembly of the appendages(26). Moreover, thanks to the recently revealed details on DA structure and organization (21, 59) it is tempting to speculate that TTBK2 might be able to target additional basal body proteins beside those included in our screen. In our initial screening, CP110 and Rab8a did not shift upon TTBK2 co-expression, hence we did not include them in the subsequent detailed analysis. However, as lack of mobility shift does not necessarily mean lack of phosphorylation, it would be premature to rule them out as TTBK2 substrates. Interestingly, two of CP110 binding partners, CEP97 and MPP9, have already been reported as TTBK2 substrates and their phosphorylation has been implicated in CP110/CEP97 complex removal from distal end of mother centriole (24, 27).

Extent of the mobility shift we observed for CEP164, CEP83, CEP89, CCDC92, Rabin8, and DVL3 suggested modification by multiple phosphates rather than by single phosphorylation, which we subsequently confirmed in MS-based experiments. Importantly, our analysis was able to detect not only majority of sites previously assigned as targets of TTBK2 activity (CEP164 sites(26), namely T1309, S1317, S1347, and S1443 in both datasets and S1346 *in vivo*), but also many new ones, hence demonstrating both robustness and sensitivity of our workflow.

Majority (64%) of TTBK2 phosphorylation sites we detected *in vitro* were subsequently confirmed *in vivo*. Nonetheless, significant portion of sites detected by *in vivo* analysis (in total 106 phosphorylated S/T residues) were not revealed using the *in vitro* approach. Several factors may account for that, such as different activity of the kinase *in vitro* and *in vivo*, different preferences for substrates or presence/absence of priming kinase(s) and counteracting phosphatases(60). TTBK2 is considered as distant member of CK1 kinase family. However, it was previously shown that its truncated variant (TTBK2 1-450) preferentially phosphorylates peptides with priming phosphotyrosine residue at position +2 (50). This unusual substrate preference, unique among CK1 family members which typically favour stretch of acidic residues or pS/pT around position −3 (55, 61–63) was related to differences in the putative phosphate-binding groove between TTBK1/2 and all remaining CK1 isoforms (50). Although several Y kinases have been proposed to reside on centrosome or mother centriole, respectively (64, 65), significance of that for centriole biogenesis or ciliogenesis is questionable. Importantly, this study as well as others (24, 26, 38, 40) failed to detect any significant presence of Y at +2 at verified TTBK2 substrates, raising a question if TTBK2 extensively utilized such motif *in vivo*. Additionally, although we detected several phosphorylations that fit the canonical consensus motif pS/pT-x-x-**S/T** of CK1, our unbiased motif search analysis implies that TTBK2 may in addition favour glutamic acid (E) on position +3, and leucine (L) at position +1, respectively, at least in a subset of its targets. These motifs resemble noncanonical CK1 motifs SLS and to some extent also a motif with acidic stretch of residues located C-terminally from the target S/T reported before (29, 66–68). Importantly, one has to bear in mind that, in contrast to motifs identified in this study and the studies where noncanonical motifs for CK1 were defined, delineation of kinase motifs has been typically carried out using rather reduced libraries of synthetic peptides, that lack secondary or tertiary structure of *bona fide* physiological substrates of a given kinase. In fact, it has been demonstrated that the phosphorylation of a substrate by CK1 isoforms does not strictly depend on the consensus sequence, but there is significant effect of tertiary structure of the substrate(69). From this point, full length TTBK2 does not seem to use the motif with pY at position +2 in substrates we examined, instead our data suggest it may use motifs resembling noncanonical motif reported for CK1. Of note, results of the assay that identified pY+2 as preferred motif for TTBK2 also show that TTBK2 can phosphorylate peptides carrying noncanonical CK1-like motifs *in vitro*, albeit to lesser extent than pY+2 (50). To further analyse the preference for motif usage by TTBK2, co-crystallization of kinase domain of TTBK2 with some of its physiological substrates would help defining the molecular mechanism mediating their mutual interactions.

The amino acid sequences of intrinsically disordered proteins determine their inability to fold into stable tertiary structures and instead enable them to rapidly interconvert between distinct conformations to mediate their biological functions (53, 70). Consequently, this allows the intrinsically disordered proteins to expose significant part of their primary sequence for binding and post-translational modification(47, 53, 54), hence provides them with ability to act as hubs in protein complexes that typically facilitate assembly of various organelles (53, 58, 71). In agreement with these reports, all proteins we identified as substrates of TTBK2 modified by multiple phosphorylations are predicted to have long IDRs suitable for such extensive modification. Given that phosphorylation of IDRs might affect proteins structural and hence binding properties (52, 58), it is tempting to speculate that TTBK2 phosphorylation allows basal body DA proteins to achieve novel quality that ensures their timely interactions and assembly to complexes necessary for the formation of PC.

We provided experimental evidence supporting this hypothesis on the example of CEP164-TTBK2 interactions. We admit that the observed effects of the introduced S to A mutations of CEP164 N-term on sequestration of TTBK2 and ciliogenesis are modest. However this is to be expected, as the mutated residues were selected while bearing in mind that they lay within IDR outside of the reported TTBK2 binding region(26), but still in the vicinity to possibly mediate effect on CEP164-TTBK2 interaction. From this perspective this data suggests that at least some of the mutated residues participate in WW domain-mediated CEP164-TTBK2 interaction. Interestingly, recent elegant work by Xu and colleagues (28) pinpointed regulatory role of a short, basic AA stretch (110-122) of CEP164 for the CEP164-TTBK2 interaction via Phosphatidylinositol-4 phosphate (PtdIns(4)P) binding. As the cluster S129-S136, phosphorylated by TTBK2, localizes very close to the described PI binding stretch, it is tempting to speculate that phosphorylation of this cluster might represent a mechanism of how TTBK2 modulates its own interaction with CEP164 via crosstalk with INPP5E/PIP3K regulatory element, plausibly by hampering binding of the acidic phosphoinositol head group to its basic binding site (28). It should be rewarding to explore whether there indeed is any functional interaction between TTBK2 sites we identified and activity of PIP3K/INPP5E implied in regulation of CEP164-TTBK2 binding (28).

Additionally, we confirmed and extended recent report on TTBK2 phosphorylation of DVL3 (51), a component of WNT signalling pathways known to reside to centrosome (48, 72, 73). This may have several biological implications, as TTBK2 seems to phosphorylate small cluster of S/T residues in the C-terminus of DVL3, which is important for regulating DVL3 oligomerization (74), which is in turn essential for DVL-mediated signal transduction via formation of signalosomes (75). In addition, the C-terminus of DVL is typically phosphorylated by CK1ε (74), thus opening an intriguing window for a crosstalk at the level of TTBK2-CK1ε, possibly through differential phosphorylation of DVL3 C-terminus as CK1ε phosphorylates DVL3 on many additional sites to those targeted by TTBK2 (51). Clearly, additional work will be necessary to fully examine the implications of TTBK2-mediated DVL phosphorylation for WNT signalling pathways as well as basal body-related aspects of DVL biology.

In summary, our findings expand current knowledge about TTBK2 substrates and provide important insight into extent and character of TTBK2-mediated phosphorylations in the context of basal body proteins and PC initiation, respectively. Underpinning the functions of TTBK2 phosphorylations as well as its interactions with other kinases implicated in ciliogenesis will undoubtedly be vital for a mechanistic understanding of cilium assembly.

## Supporting information

Supplementary figures as referred to in the manuscript text

Supplementary table 1 (phosphosite mapping)

Supplementary table 2 (kinase motif delineation)

Supplementary table 3 (plasmids, antibodies list)

## Acknowledgement

We thank Erich Nigg, Gislene Pereira, Vitezslav Bryja, Peter Jackson, Robert Lefkowitz, Stephane Angers, Joon Kim, and Lumir Krejci for sharing reagents or instruments. The work was supported by grants from Czech Science Foundation (16-03269Y, 19-05244S) and Swiss National Science Foundation (IZ11Z0_166533) to LC. OB was supported by funds from the Faculty of Medicine MU to junior researcher (Ondrej Bernatik). KH and ZZ were supported by the project CEITEC 2020 (LQ1601) funded by the Ministry of Education, Youth and Sports of the Czech Republic (MEYS). We acknowledge the core facility CELLIM of CEITEC, supported by the MEYS CR (LM2015062 Czech-Bio-Imaging). CIISB research infrastructure project LM2015043 funded by MEYS CR is gratefully acknowledged for the financial support of the LC-MS/MS measurements at the Proteomics Core Facility.

## FOOTNOTES

### Abbreviations used are

TTBK2: Tau tubulin kinase 2;
KD: kinase dead;
PC: Primary cilium;
DA: Distal appendages;
SDA: sub-distal appendages;
CK1: casein kinase 1;
SCA11: spinocerebellar ataxia type 11;
PONDR: Predictor of Natural Disordered Regions;
IDRs: intrinsically disordered regions;
DEPP: Disorder Enhanced Phosphorylation Predictor;
CEP164 N-term: N-terminal part of CEP164;
PtdIns(4)P: phosphatidylinositol (4) phosphate;

