## Supplementary figures as referred to in the manuscript text for "Phosphorylation of multiple proteins involved in ciliogenesis by Tau Tubulin kinase 2"

### Supplementary figure 1

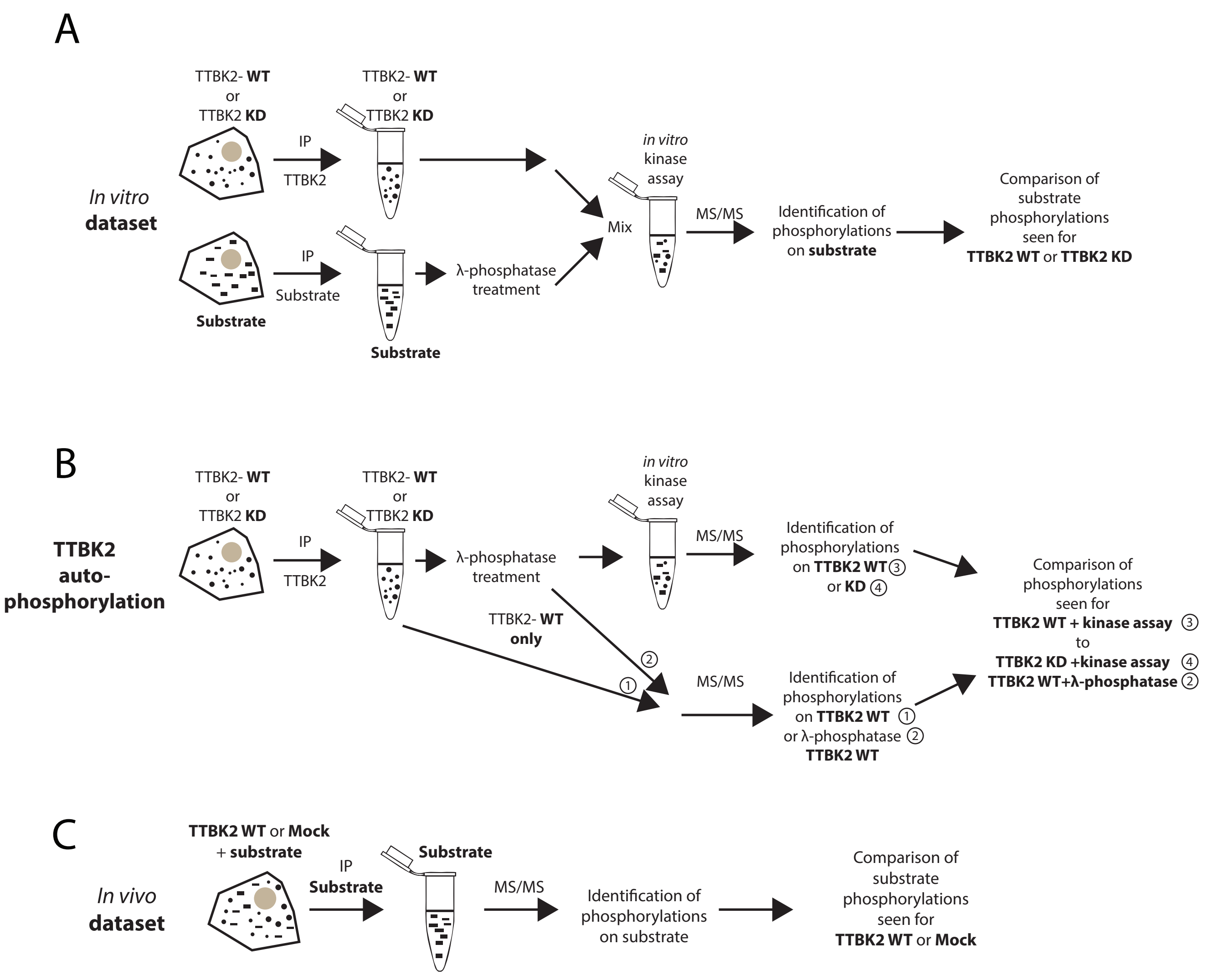


**Supplementary figure 1. Schematized workflow of MS/MS experiments**
**A:** HEK293T cells were transfected by tested substrate or TTBK2 WT/KD. Proteins of interest were purified by immunoprecipitation. Additional treatment with λ phosphatase was performed in second experimental repeat. Tested substrates proteins were subjected to in vitro kinase assay with TTBK2 WT or KD (control). Following SDS-PAGE separation samples were analysed by MS/MS to identify phosphorylations. **B: T**ransfected TTBK2 WT or TTBK2 KD were isolated from HEK293T cells by immunoprecipitation. In total, four conditions were prepared, as indicated **-** ① TTBK2 WT**,** ② TTBK2 WT treated with λ-phosphatase, ③ TTBK2 WT treated with λ-phosphatase subsequently subjected to in vitro kinase assay, and ④ phosphatase-treated TTBK2 KD subjected to in vitro kinase assay. Following SDS-PAGE separation samples were analysed by MS/MS to identify phosphorylations. Condition ③ was compared to conditions ② and ④**,** respectively based on criteria described in M&M section. Sample ① was used as reference to monitor phosphatase treatment effectivity. **C:** Tested substrate transiently expressed in HEK293T cells together with exogenous TTBK2 WT or empty plasmid (mock) was purified by immunoprecipitation and following SDS-PAGE separation analysed by MS/MS. Identified phosphorylations found in condition with TTBK2 WT were compared to those found in mock condition, based on criteria described in the M&M section.

### Supplementary figure 2

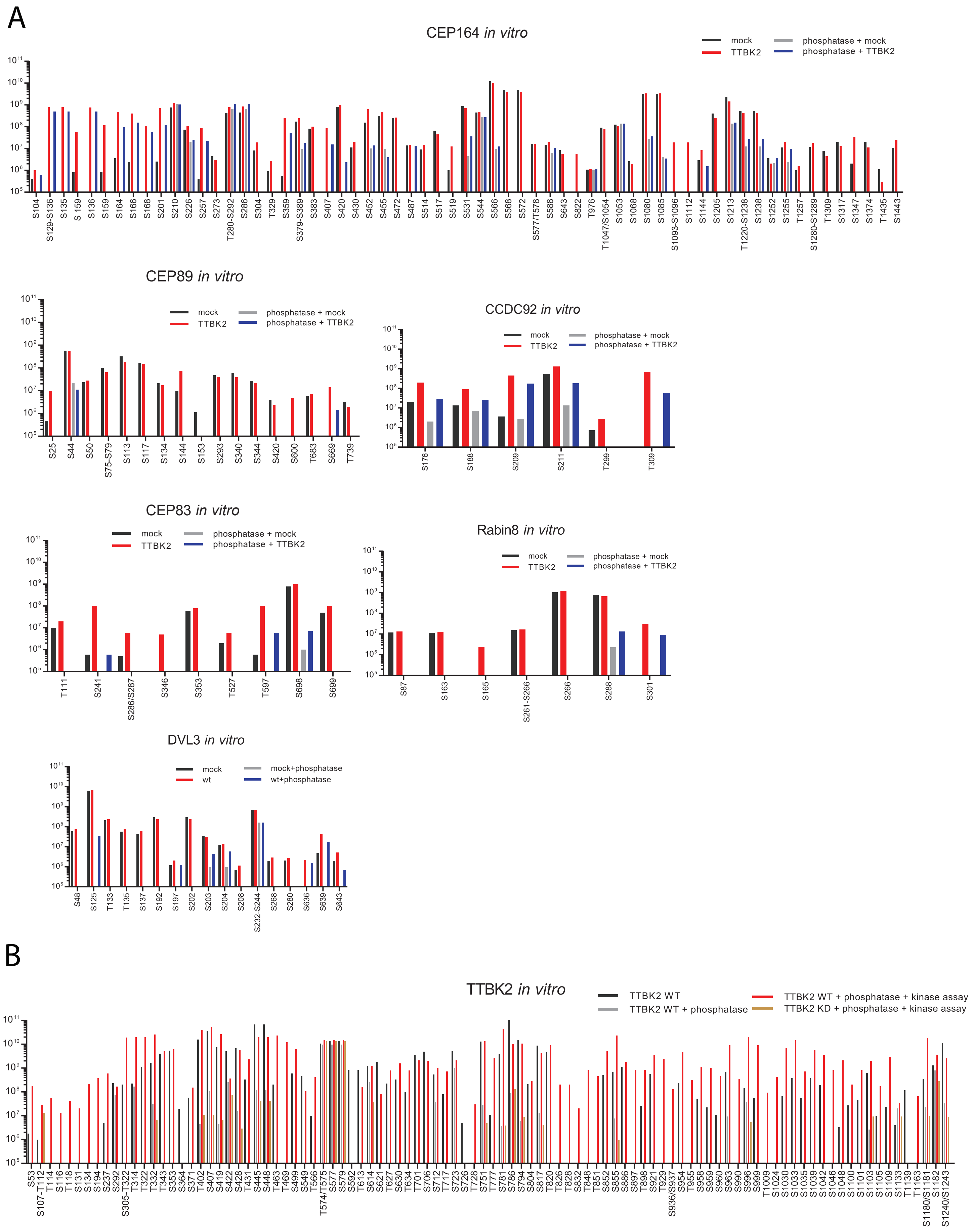


**Supplementary figure 2. Summary of phosphorylations identified for depicted substrates *in vitro*.**
For each S/T site on x axis, average phosphorylation intensity is shown in y axis (log10 scale) for indicated conditions. Cases where the exact position could not be assigned are indicated as (/) for two possible positions or (-) for clusters. **A:** Graphical summary of phosphorylations detected in vitro in following conditions. Black – Substrate + TTBK2 KD (Mock), Red – Substrate + TTBK2 WT, Grey – Substrate + λ phosphatase treatment + TTBK2 KD (Mock), Blue – Substrate + λ phosphatase treatment + TTBK2 WT. **B:** Graphical summary of phosphorylations detected in TTBK2 autophosphorylation dataset in following conditions : ① Black –TTBK2 WT, ② Grey – TTBK2 WT+ λ phosphatase treatment (Ctrl#1), ③ Red – TTBK2 WT + λ phosphatase treatment + kinase assay, ④ Brown – TTBK2 KD + phosphatase treatment + kinase assay (Ctrl#2).

### Supplementary figure 3

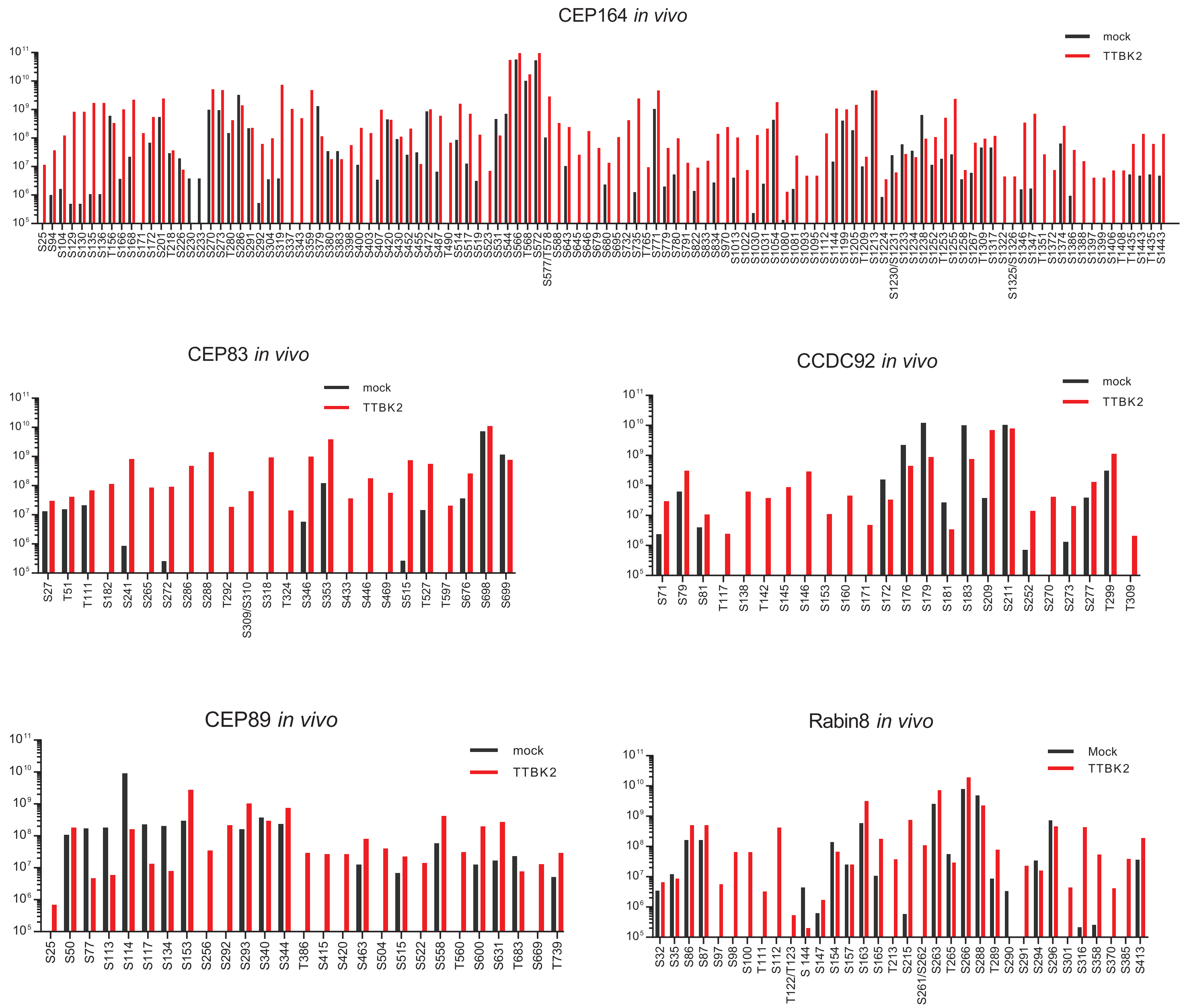


**Supplementary figure 3. Summary of phosphorylations identified for depicted substrates *in vivo*.**
For each S/T site on x axis, average phosphorylation intensity is shown in y axis (log10 scale) for indicated conditions. Cases where the exact position could not be assigned are indicated as (/). Graphical summary of phosphorylations detected in vivo in following conditions: Black – Substrate + Mock, Red – Substrate + TTBK2.

### Supplementary figure 4

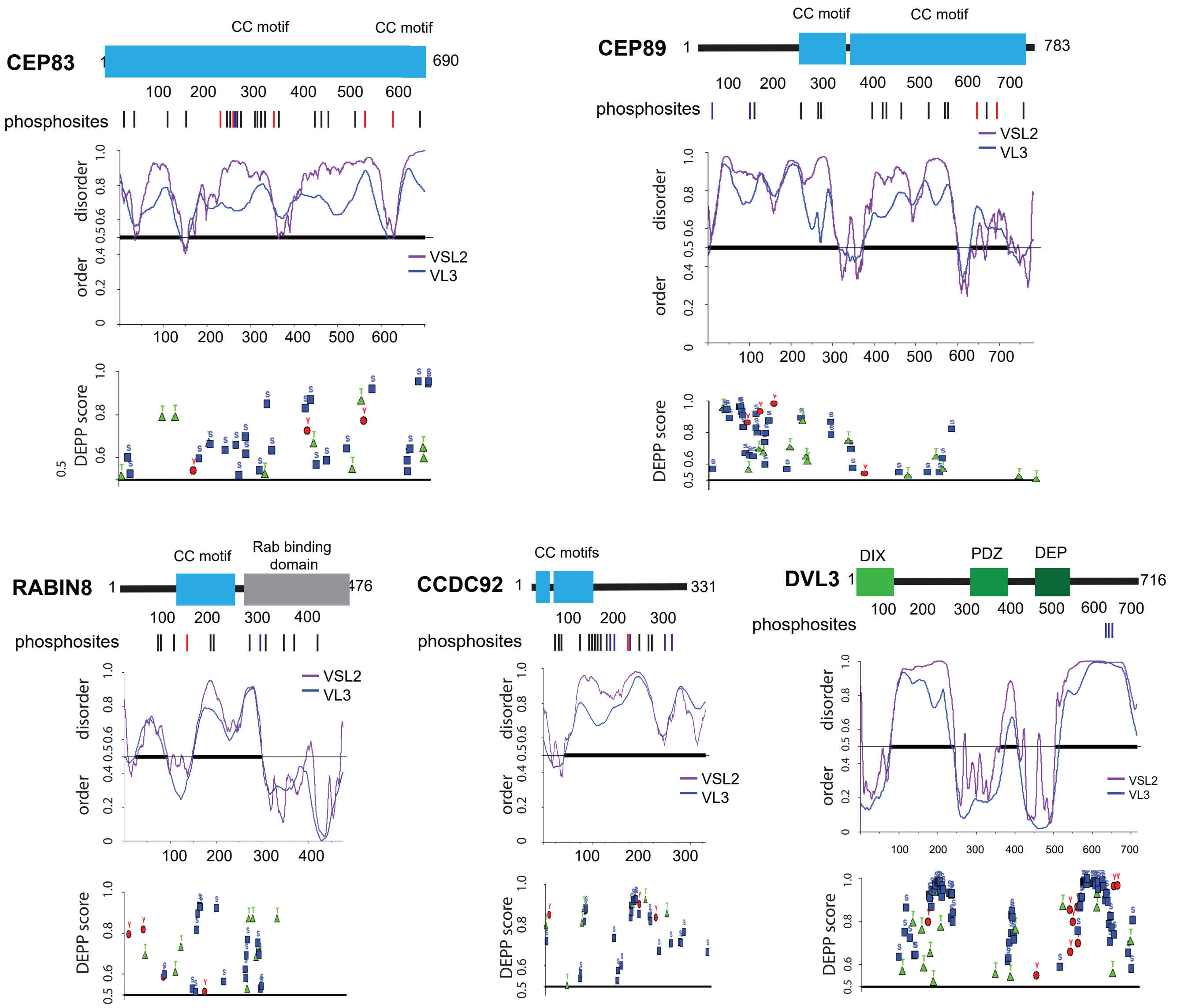


**Supplementary figure 4. Disorder predictions of CEP89, CEP83, CCDC92, Rabin8 and DVL3 indicate long IDRs with multiple phosphorylation sites.**

Protein disorder and phosphorylation prediction for CEP89, CEP83, CCDC92, Rabin8 and DVL3. Structure of all proteins is schematized, rectangles indicate presence of a domain or motif, the numbers indicate length of given protein in amino acids. Lines below the schematic protein structure indicate positions of each phosphorylation induced by TTBK2. The first graph below the schematized protein shows prediction of order/disorder (with cut-off 0.5) determined by PONDR using VSL2 (Violet line) and VL3-BA (Blue line) algorithms. The second graph plots Disorder Enhanced Phosphorylation Predictor (DEPP) score (with cut-off score 0.5) and individual phosphosites thereby predicted (pS as blue rectangle, pT as green triangle, and phosphoY as red circle).

### Supplementary figure 5

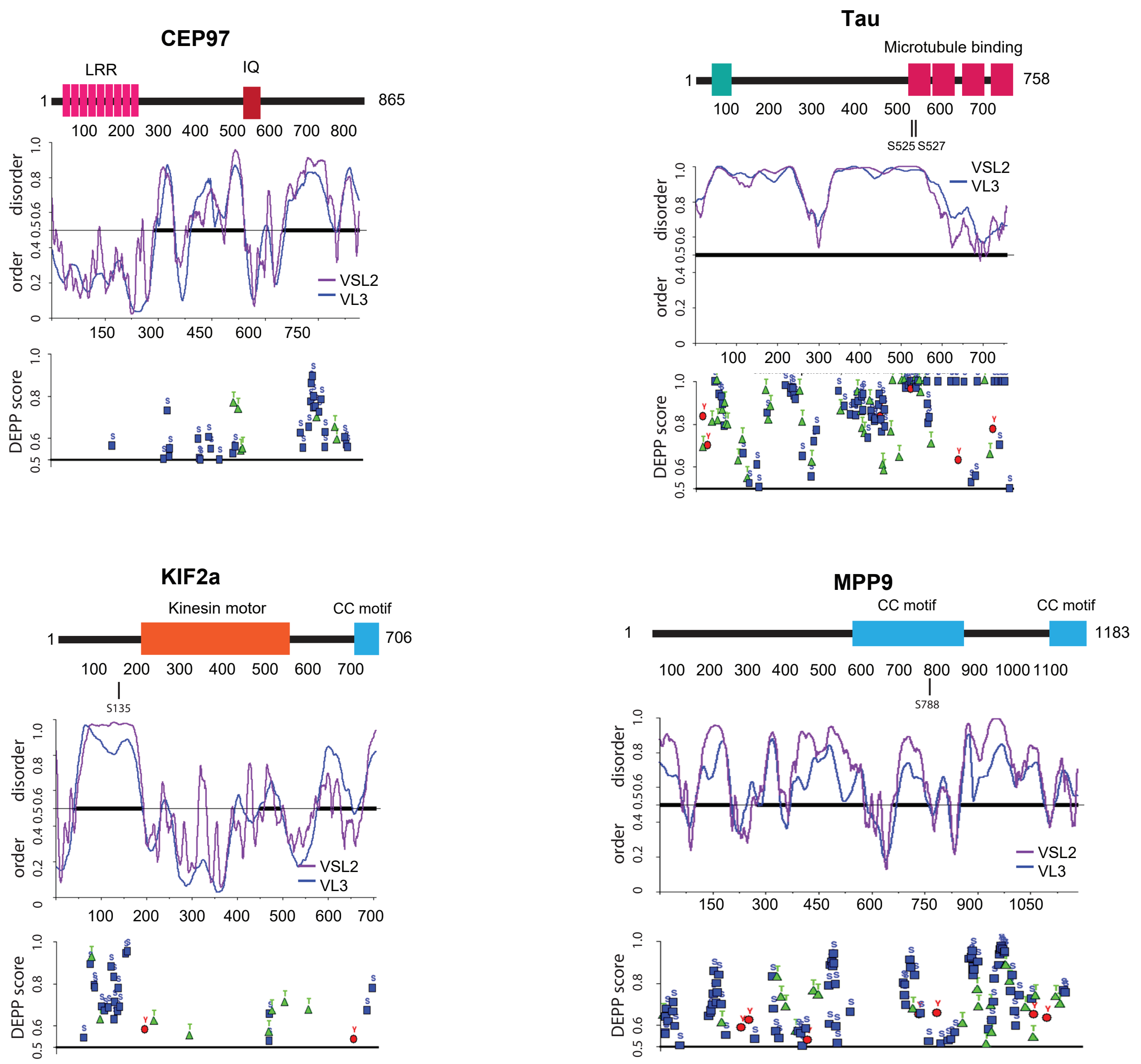


**Supplementary figure 5. Disorder predictions of CEP97, KIF2a, Tau and MPP9 indicate long IDRs with multiple phosphorylation sites.**
Protein disorder and phosphorylation prediction for CEP97, KIF2a, MPP9 and Tau proteins. Structure of all proteins is schematized, rectangles indicate presence of a domain or motif, the numbers indicate length of given protein in amino acids. Lines and numbers respectively, below protein structure indicate published TTBK2 phosphorylated residues (24, 29, 38, 40). The first graph below the schematized protein shows prediction of order/disorder (with cut-off 0.5) determined by PONDR using VSL2 (Violet line) and VL3-BA (Blue line) algorithms. The second graph plots Disorder Enhanced Phosphorylation Predictor (DEPP) score (with cut-off score 0.5) and individual phosphosites thereby predicted (pS as blue rectangle, pT as green triangle, and phosphoY as red circle).
